# ERK1/2-dependent activation of FCHSD2 drives cancer cell-selective regulation of clathrin-mediated endocytosis

**DOI:** 10.1101/346593

**Authors:** Guan-Yu Xiao, Aparna Mohanakrishnan, Sandra L. Schmid

## Abstract

Clathrin-mediated endocytosis (CME) regulates the uptake of cell surface receptors, as well as their downstream signaling activities. We recently reported that signaling reciprocally regulates CME in cancer cells and that the crosstalk can contribute to cancer progression. To further explore the nature and extent of the crosstalk between signaling and CME in cancer cell biology, we analyzed a panel of oncogenic signaling kinase inhibitors for their effects on CME. Inhibition of several kinases selectively affected CME function in cancer cells. Among these, ERK1/2 inhibition selectively inhibited CME in cancer cells by decreasing the rate of CCP initiation. We identified an ERK1/2 substrate, the FCH/F-BAR and SH3 domain-containing protein, FCHSD2, as being essential for the ERK1/2-dependent effects on CME and CCP initiation. ERK1/2 phosphorylation activates FCHSD2 and regulates EGFR endocytic trafficking as well as downstream signaling activities. Loss of FCHSD2 activity in non-small-cell lung cancer cells leads to increased cell surface expression and altered signaling downstream of EGFR, resulting in enhanced cell proliferation and migration. The expression level of FCHSD2 is positively correlated with higher cancer patient survival rate, suggesting that FCHSD2 negatively affects cancer progression. These findings provide new insight into the mechanisms and consequences of the reciprocal regulation of signaling and CME in cancer cells.

**Significance:** Clathrin-mediated endocytosis (CME) determines the internalization of receptors and their downstream signaling. We discovered that CME is differentially regulated by specific signaling kinases in cancer cells. In particular, ERK1/2-mediated phosphorylation of the FCH/F-BAR and double SH3 domains-containing protein 2 (FCHSD2) regulates CME, and the trafficking and signaling activities of EGF receptors. This reciprocal interaction negatively regulates cancer proliferation and migration. The expression level of FCHSD2 is positively correlated with higher cancer patient survival rates. This study identifies signaling pathways that impinge on the endocytic machinery and reveals a molecular nexus for crosstalk between intracellular signaling and CME. Cancer cells specifically adapt this crosstalk as a determinant for tumor progression, which has implications for novel therapeutics against cancers.

## Introduction

Cancer progression involves tumor cell proliferation and metastasis driven, in part, by altered intracellular signaling downstream of plasma membrane receptors (1, 2). In tumor cells, signaling activities can be altered by dysregulated endocytic trafficking that increases receptor recycling and decreases lysosomal targeting and degradation (3-5). Hence, a link between endocytosis and cancer behavior has been suggested (3, 6-8). Clathrin-mediated endocytosis (CME) is the major endocytic pathway that determines the rates of internalization of plasma membrane receptors, regulates their cell surface expression, and controls their downstream signaling activities (9, 10). Despite the well-known fact that CME can regulate intracellular signaling pathways (11), whether and how signaling can feedback to regulate CME during cancer cell progression has been less studied.

CME occurs when clathrin-coated pits (CCPs) assemble on the cell surface, concentrate cargo molecules (i.e. receptors and their bound ligands), invaginate and undergo membrane fission catalyzed by the GTPase dynamin to release clathrin coated vesicles into the cytosol. Once considered a constitutive process, recent studies have demonstrated that CME can be dynamically regulated in response to cargo and downstream signaling pathways (12-15). For example, in non-small cell lung cancer (NSCLC) cells, dynamin-1 (Dyn1), the neuron-enriched isoform, is frequently overexpressed and can be activated downstream of the Akt/GSK3β signaling cascade, resulting in increased rates of CCP initiation and dysregulated CCP maturation (15). Moreover, the clathrin light chain isoform b (CLCb), which functions primarily to regulate clathrin assembly and is also neuronally-enriched (16, 17), is specifically upregulated in NSCLC cells (18). CLCb-dependent alterations in CME lead to Dyn1 activation via increased Akt/GSK3β phosphorylation associated with APPL1-positive signaling endosomes (18). The crosstalk between signaling and CME contributes to abnormal trafficking of epidermal growth factor receptors (EGFRs) and altered downstream signaling, leading to enhanced cancer cell migration and metastasis (18). Accordingly, it is important to further understand the nature and extent of interactions between CME and intracellular signaling in cancer cell biology.

To address this issue, we analyzed a panel of oncogenic signaling kinase inhibitors for their differential effects on CME in several human non-cancerous and cancer cells. We identified ERK1/2 activity as being selectively required for rapid CME and CCP initiation in cancer cells and FCHSD2 (Nervous Wreck, in *Drosophila*) as the downstream kinase substrate responsible these effects. Our study provides new insight into the mechanisms and consequences of the reciprocal regulation of signaling and CME in cancer cells.

## Results

### Oncogenic Signaling Differentially Affects CME in Cancer Cells

Recent studies have shown that the crosstalk between intracellular signaling and CME can affect cancer progression and metastasis (8, 18). Thus, it is important to determine the extent of this crosstalk as well as the signaling pathways that impinge on the regulation of CME in cancer cells. To this end, we prioritized a set of kinases and GTPases that are often implicated in cancer-relevant signaling pathways and assessed the effects of their validated inhibitors (19) on transferrin receptor (TfnR) endocytosis, a canonical marker for the quantification of CME (20, 21). Initially, we chose non-cancerous ARPE-19 cells, which have been routinely used to study TfnR endocytosis (20) and compared these to H1299 NSCLC cells, which exhibit crosstalk between signaling and CME (15, 18).

Strikingly, 10 of the 21 inhibitors we screened differentially affected CME in H1299 vs ARPE cells (Figure S1). This included inhibition of Akt, which we previously showed functions through an Akt/GSK3β signaling cascade to activate Dyn1 and enhance endocytosis in H1299 and A549 NSCLC cells (15, 18, 22). To further validate these findings, we extended the functional comparison of these 10 inhibitors to more closely matched cancerous and non-cancer cell lines. We used another NSCLC cell line, HCC4017, together with the syngeneic non-tumorigenic HBEC line, HBEC30KT, derived from the same patient (23). In addition, we compared the breast cancer cell line, MDA-MB-231 with normal human mammary epithelial cells, MCF10A (24). Consistent with our initial screen, inhibition of a subset of these kinases significantly reduced CME activity in most cancer cell lines, without affecting CME in their non-cancerous counterparts (Figure 1A). These findings reveal the unexpected differential regulation of CME in cancer cells by distinct intracellular signaling pathways.

**Figure 1.**
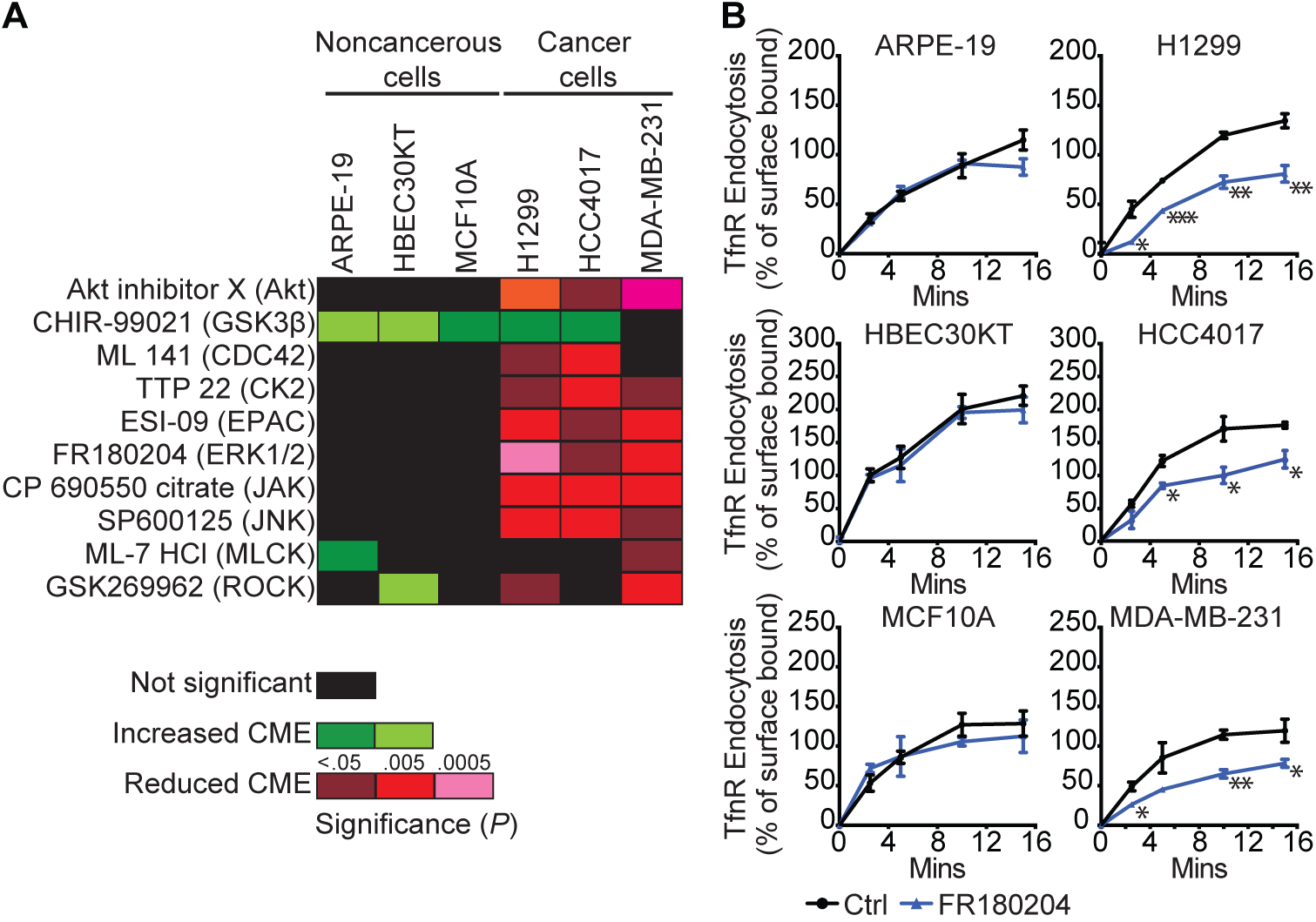
Differential effects of kinase inhibitors on CME in cancer cells. (*A*) Heatmap illustrating results from a systematic analysis of the effects of pharmacological kinase inhibitors on TfnR endocytosis in human non-cancerous and cancer cell lines. (*B*) Endocytosis of TfnR was measured in different pairs of human non-cancerous and cancer cells without or with the ERK1/2 inhibitor (FR180204, 10 µM) treatment. Percentage of internalized TfnR was calculated relative to the initial surface TfnR. Data represent mean ± SEM (*n* = 3). Two-tailed Student’s *t* tests were used to assess statistical significance for comparison with Ctrl. \**P* < 0.05, \*\**P* < 0.005, \*\*\**P* < 0.0005.

Inhibiting some of these kinases selectively enhanced CME in non-cancerous vs. cancer cells. GSK3β enhanced rates of CME in most cell lines tested, presumably through activation of Dyn1 (15, 22). Interestingly, inhibition of myosin light chain kinase (MLCK) and Rho kinase (ROCK) accelerated endocytosis in a subset of non-cancer cell lines, while inhibiting CME in a subset of cancer cells. As these kinases control the actin cytoskeleton, these results could reflect known cell-type differences in the role of actin in cancer vs non-cancer cells (25, 26). While it will be important to investigate how each of these signaling pathways impinge on CME, we chose to focus our further studies on ERK signaling as the EGFR/Ras/ERK1/2 pathway is a major driver of, and therapeutic target for multiple cancers (27-30). Inhibition of ERK1/2 selectively inhibits CME in all cancer cell lines tested without affecting CME in several non-cancerous cell lines (Figure 1B). For further studies we chose ARPE-19 and H1299 cells, as well as the syngeneic pair, HCC4017 and HBEC30KT.

### ERK1/2 Specifically Regulates CME in Cancer Cells

ERK1/2 signaling, downstream of EGFR and Ras, is highly activated in many cancers. To confirm that ERK1/2 activity is selectively involved in the regulation of CME in cancer cells we used a second ERK1/2 inhibitor (SCH772984), as well as a well-characterized inhibitor of the essential upstream kinase, MEK (GSK1120212). As expected, these ERK1/2 signaling inhibitors efficiently decreased ERK1/2 phosphorylation (Figure 2A and S2A). Importantly, they also specifically inhibited CME both in H1229 relative to ARPE-19 cells (Figure 2B) and in HCC4017 relative to HBEC30KT cells (Figure S2B).

**Figure 2.**
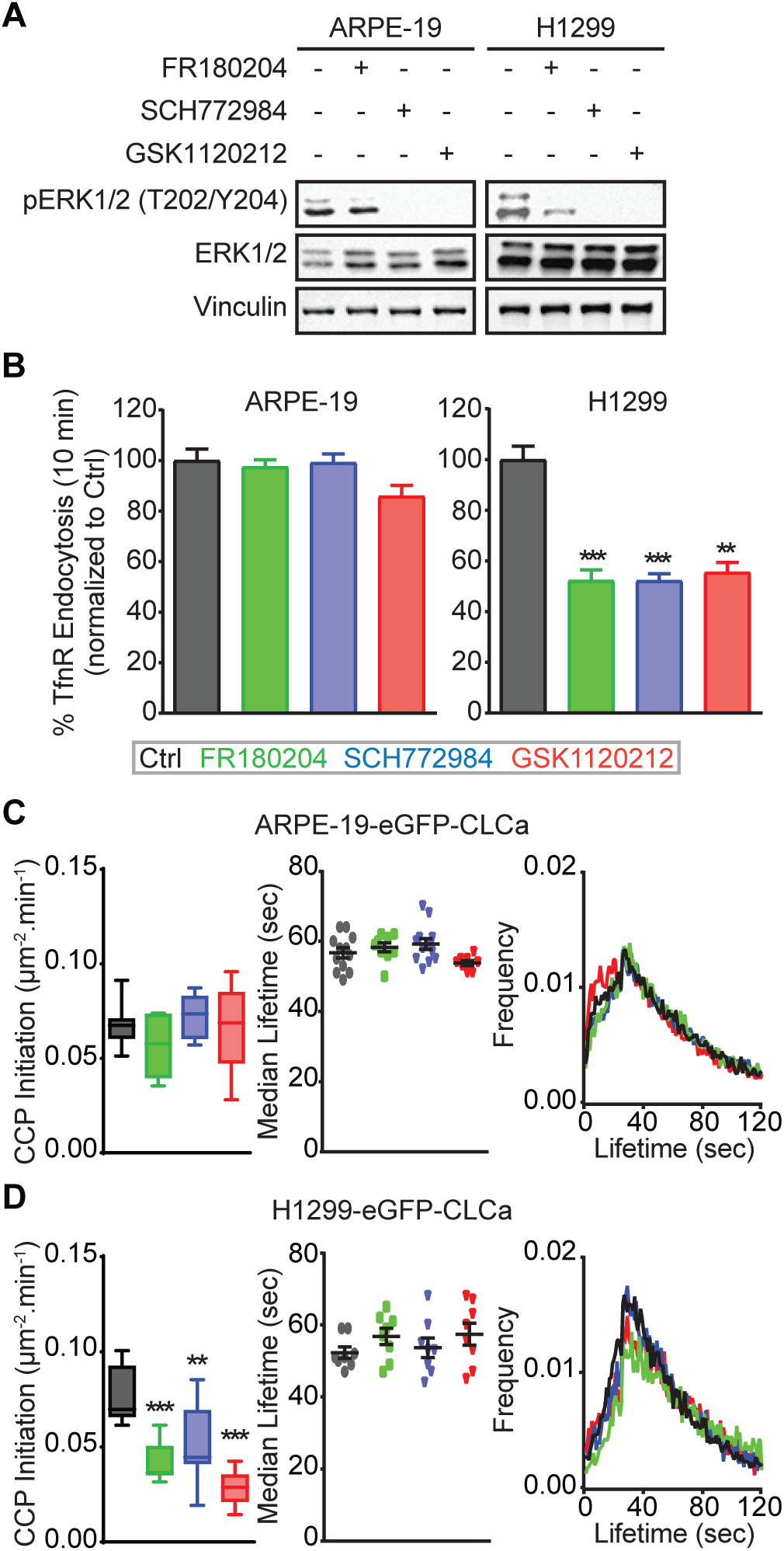
ERK1/2 selectively affects CME activities in cancer cells. (*A*) Representative western blots used to measure the efficiencies of kinase inhibitors in reducing ERK1/2 phosphorylation in control, ERK1/2 inhibitor (FR180204 and SCH772984, 10 µM) and MEK1/2 inhibitor (GSK1120212, 10 µM) treated ARPE-19 and H1299 cells. (*B*) Endocytosis of TfnR was measured in ARPE-19 and H1299 cells untreated (Control) or incubated with the indicated inhibitors, as described above. All data were normalized to the % of TfnR internalized after 10 min in control cells and represent mean ± SEM (*n* = 3). Average lifetime distributions (*left*), and rates of initiation of *bona fide* CCPs in control and inhibitor-treated (*C)* ARPE-19 cells and (*D*) H1299 cells. Data in (*C*) and (*D*) were obtained from at least 15 cells/condition, the box plots represent median, 25th and 75th percentiles, and outermost data points. Two-tailed Student’s *t* tests were used to assess statistical significance for comparison with Ctrl. \*\**P* < 0.005, \*\*\**P* < 0.0005.

CME is a multistep process that involves CCP initiation, stabilization, maturation and finally fission. To determine which step(s) of CME were affected upon ERK1/2 inhibition, we analyzed CCP dynamics using quantitative total internal reflection fluorescence microscopy (TIRFM) (15, 18) in ARPE-19 and H1299 cells stably expressing EGFP-clathrin light chain a (EGFP-CLCa), and in HCC4017 cells stably expressing SNAP-CLCa. As expected, CCP dynamics were unchanged in the ARPE-19 cells upon inhibitor treatment (Figure 2C). However, inhibition of ERK1/2 signaling dramatically reduced the rate of CCP initiation without significantly altering the lifetime distribution or median lifetimes of CCPs in both H1299 cells (Figure 2D) and HCC4017 cancer cells (Figure S2C). These data indicate that ERK1/2 signaling differentially regulates the first key step of CME (i.e. CCP initiation) (31) in cancer cells.

### FCHSD2 Regulates CME in Cancer Cells

ERK1/2 kinases are ubiquitously expressed serine/threonine kinases that regulate diverse biological processes (32-35) by phosphorylating multiple substrates (36). We screened the literature for ERK1/2 substrates and identified FCHSD2 (37), the human orthologue of Nervous Wreck (Nwk) in *Drosophila*, which has been implicated in membrane remodeling (38) and the regulation of growth factor signaling (39, 40) at the synapse. While thus far not studied in the context of CME in mammalian cells, FCHSD2 was of interest, because it encodes an N-terminal F-BAR domain predicted to initiate membrane curvature and two C-terminal SH3 domains proposed to interact with the components of the actin and CME machinery (41).

To test its potential role in CME, we first investigated the effects of efficient siRNA-mediated knockdown of FCHSD2 (Figure 3A and S3A) in non-cancerous and cancer cells. Like ERK1/2 inhibition, depletion of FCHSD2 did not significantly alter CME activities in non-tumorigenic ARPE-19 or HBEC30KT cells, but potently inhibited CME in H1299 and HCC4017 NSCLC cells (Figure 3B and S3B). FCHSD2 depleted cancer cells were no longer sensitive to ERK1/2 inhibition (Figure 3C and S3C), even though they remained sensitive to SP600125 (Figure S3C), a JNK inhibitor that also specifically affects CME in cancer cells (see Fig. 1A). These data suggest that FCHSD2 might be the CME-relevant downstream target of ERK. Consistent with this, like ERK inhibition, knockdown of FCHSD2 specifically inhibited the rate of CCP initiation without affecting CCP lifetimes (Figure 3D). The reduced rate of CCP initiation in FCHSD2- depleted cells was now insensitive to ERK1/2 inhibition. Together, these data demonstrate that FCHSD2 plays a role in CME, especially in CCP initiation, and suggest that it functions as a relevant downstream effector of ERK1/2 kinases.

**Figure 3.**
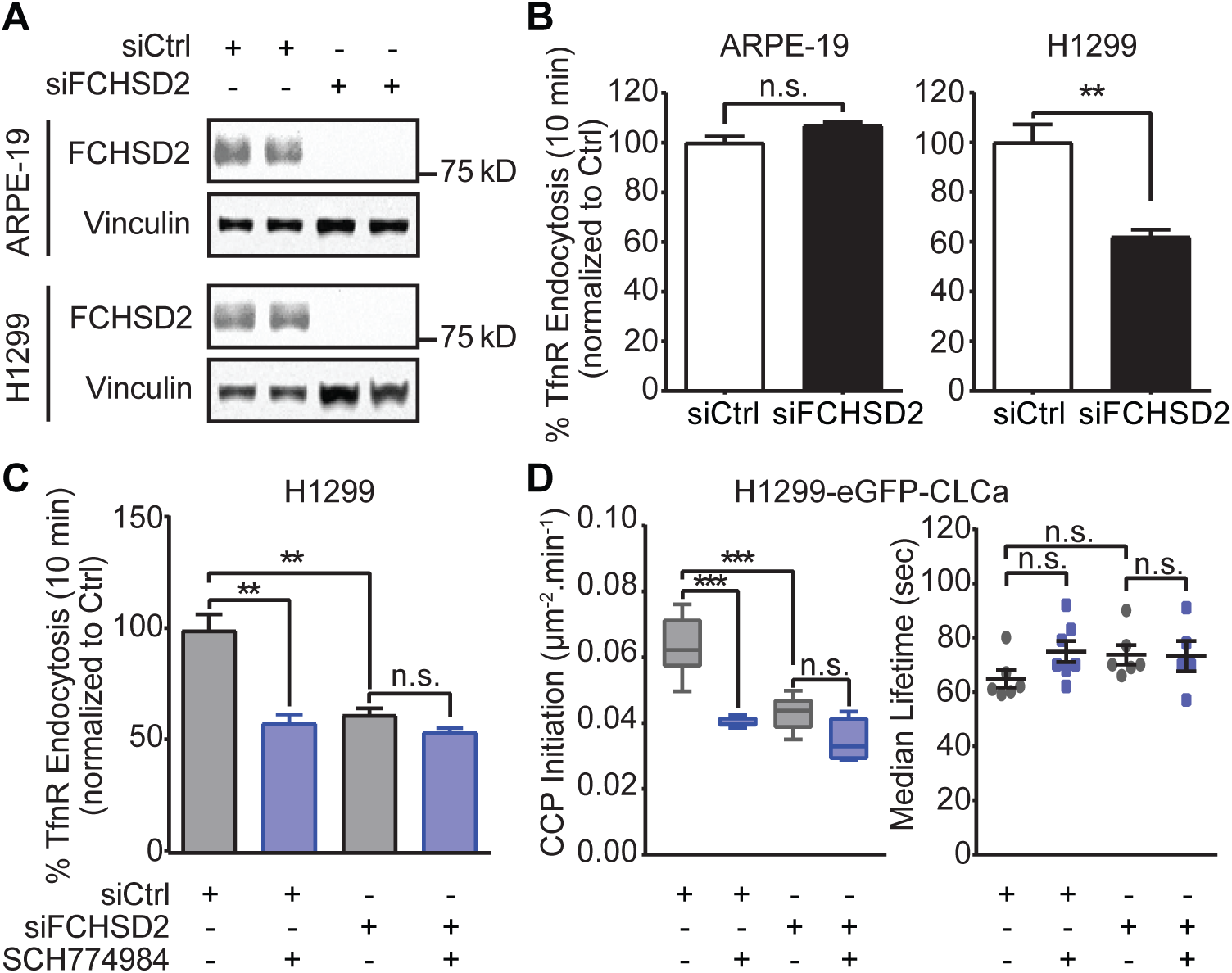
FCHSD2 is specifically enlisted for CME in cancer cells. (*A*) Representative western blots and the quantification of FCHSD2 knockdown efficiency in control and FCHSD2 siRNA-treated ARPE-19 and H1299 cells. (*B*) Endocytosis of TfnR was measured in control and FCHSD2 siRNA-treated ARPE-19 and H1299 cells. All data were normalized to control as in Fig. 2. (*C*) Endocytosis of TfnR were measured, as described in (*B*), in control and FCHSD2 siRNA-treated H1299 cells without or with the ERK1/2 inhibitor (SCH772984, 10 µM) treatment. Data in (A)-(*C*) represent mean ± SEM (*n* = 3). (*D*) Average lifetime distributions (*left*), and rates of initiation (# CCP/µm^2^/min, *right*) of *bona fide* CCPs in control and FCHSD2 siRNA-treated H1299 cells without or with the ERK1/2 inhibitor (SCH772984, 10 µM) treatment. Data were obtained from at least 15 cells/condition, the box plots represent median, 25th and 75th percentiles, and outermost data points. Two-tailed Student’s *t* tests were used to assess statistical significance. n.s., not significant, \*\**P* < 0.005, \*\*\**P* < 0.0005.

### ERK1/2 Directly Regulates FCHSD2-Mediated CME in Cancer Cells

The phosphorylation of FCHSD2 on S681 within a canonical ERK phosphorylation motif (PXSP) was identified in a large-scale screen for ERK2 substrates using an analogue-sensitive ERK2 mutant (37). We first confirmed phosphorylation of this site in EGF-treated HCC4017 cells by phospho-proteomic analysis (Supplemental Table 1). To test whether FCHSD2 was indeed the ERK1/2 substrate responsible cancer cell-specific regulation of CME, we performed FCHSD2 siRNA-mediated knockdown in H1299 and HCC4017 cells followed by reconstitution with either wild-type FCHSD2 (FCHSD2^WT^-Myc), a mutant that cannot be phosphorylated by ERK1/2 (FCHSD2^S681A^-Myc), or with a phosphomimetic mutant (FCHSD2^S681E^-Myc) at near endogenous levels (Figure 4A). Reconstitution of FCHSD2 siRNA-depleted H1299 or HCC4017 cells with FCHSD2^WT^-Myc rescued CME activity, which was again sensitive to ERK1/2 inhibition (Figure 4B, dark gray bars). In contrast, reconstitution with non-phosphorylatable FCHSD2^S681A^-Myc failed to restore CME or sensitivity to ERK1/2 inhibition (Figure 4B, black bars). Strikingly, reconstitution with FCHSD2^S681E^-Myc resulted in rates of CME higher than in control cells, and importantly resistance to ERK1/2 inhibition (Figure 4B, twill bars). Together these data provide strong evidence that phosphorylation of FCHSD2 is indeed required for the ERK1/2-dependent effects on CME in cancer cells.

**Figure 4.**
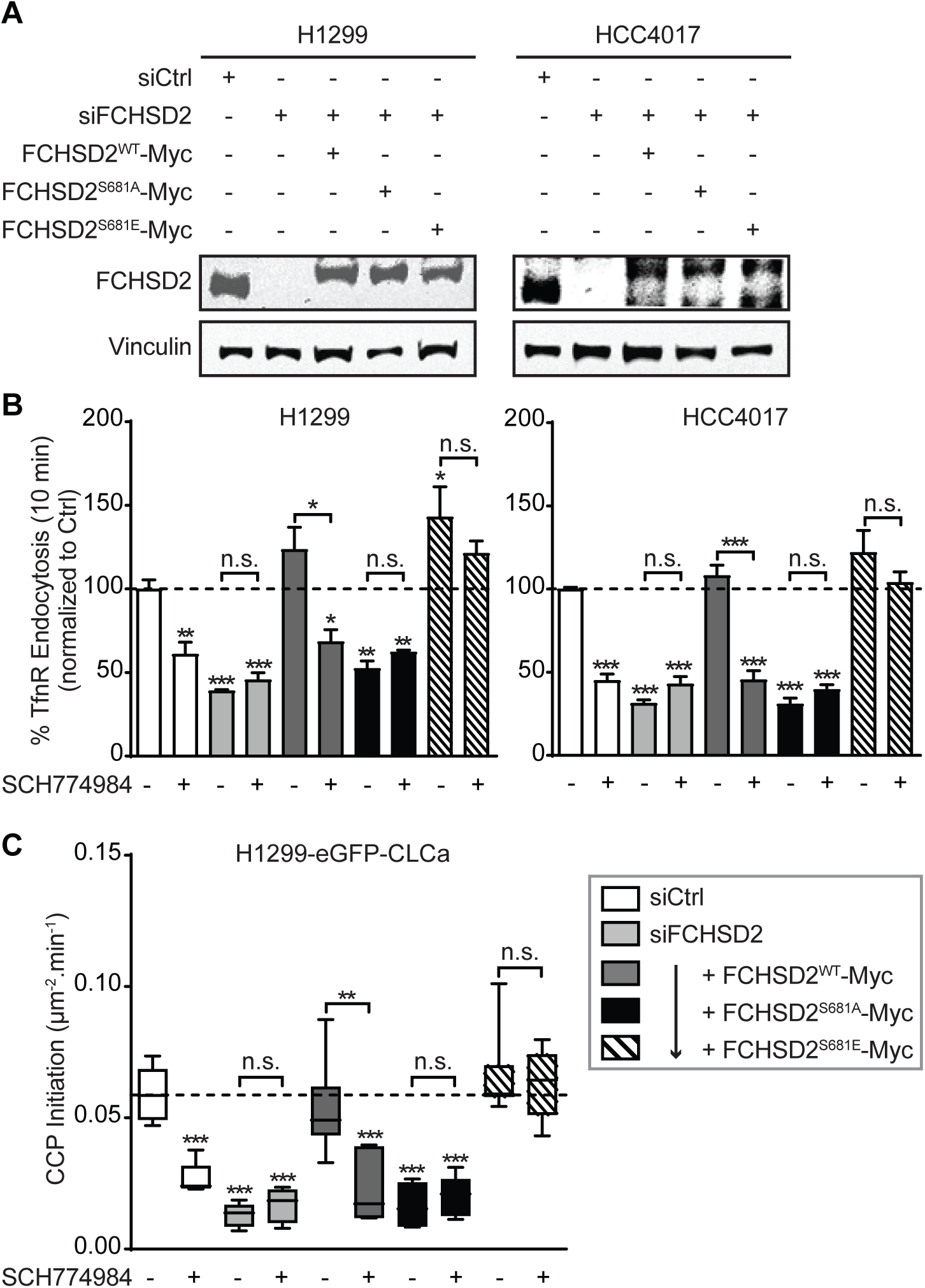
ERK1/2-mediated phosphorylation of FCHSD2 is required for CME in cancer cells. (*A*) Representative western blots of FCHSD2 expression in the control, FCHSD2 knockdown (KD) cells, or FCHSD2 KD cells reconstituted with FCHSD2^WT^-Myc, FCHSD2^S681A^-Myc, or FCHSD2^S681E^-Myc. (*B*) Endocytosis of TfnR was measured in the control, FCHSD2 KD cells, or FCHSD2 KD cells reconstituted with FCHSD2^WT^-Myc, FCHSD2^S681A^-Myc, or FCHSD2^S681E^Myc, in the absence or presence of the ERK1/2 inhibitor (SCH772984, 10 µM). All data were normalized to control and represent mean ± SEM (*n* = 3). (*C*) Rates of initiation (# CCP/µm^2^/min) of *bona fide* CCPs in the cells as described in (*B*). Data were obtained from at least 15 cells/condition, the box plots represent median, 25th and 75th percentiles, and outermost data points. Two-tailed Student’s *t* tests were used to assess statistical significance for comparison with Ctrl and for the indicated dataset. n. s., not significant, \**P* < 0. 05, \*\**P* < 0.005, \*\*\**P* < 0.0005.

Reconstitution of FCHSD2-depleted cells with FCHSD2^WT^-Myc, but not the FCHSD2^S681A^-Myc mutant, restored ERK1/2 dependent rates of CCP initiation (Figure 4C, dark gray and black bars). CCP initiation in siRNA-treated cells reconstituted with FCHSD2^S681E^-Myc were no longer responsive to ERK1/2 inhibition (Figure 4C, twill bars). These results establish that the effects of ERK1/2 inhibition on CME and CCP initiation are primarily mediated by FCHSD2 activity through phosphorylation of S681.

### FCHSD2 Regulates the Endocytic Trafficking and Surface Expression of EGFR

Many properties of progressive cancer cells are driven by altered signaling downstream of membrane receptors (42). EGFR is the predominant oncogenic receptor in NSCLC and its downstream signaling can be changed by alterations in endocytic trafficking (43, 44). Given that the crosstalk between ERK1/2 signaling and FCHSD2 functions to regulate CME, we next examined whether this crosstalk might selectively affect EGFR endocytosis in non-cancerous cells or only be active in cancer cells. Consistent with TfnR endocytosis, neither ERK1/2 inhibition nor siRNA-mediated knockdown of FCHSD2 significantly affected EGFR endocytosis in non-tumorigenic ARPE-19 cells (Figure 5A). However, EGFR uptake in HBEC30KT cells was more sensitive to ERK1/2 inhibition and FCHSD2 knockdown than TfnR uptake (compare Fig. 5A with Fig. S2B and S3B). These cell type differences might reflect the immortalization process (45) or the strong dependence on supplemental EGF for growth and maintenance of the HBEC cell line (see Methods). Alternatively, they may reflect cell-type and/or cargo-dependent differences in endocytic machinery as previously observed for death receptor-dependent endocytosis (46).

As expected, ERK1/2 inhibition reduced EGFR endocytosis in H1299 and HCC4017 NSCLC cells (Figure 5B), resulting in an ∼2-fold increase in the level of surface EGFR (Figure 5C, white bars). Knockdown of FCHSD2 similarly resulted in a significant decrease in the rate of EGFR endocytosis and an increase in surface EGFR, and these effects were insensitive to ERK1/2 inhibition (Figure 5B and C, light gray bars). Reconstitution of FCHSD2-depleted cells with FCHSD2^WT^-Myc restored EGFR endocytosis rates and EGFR surface levels, as well as sensitivity to ERK1/2 inhibition (Figure 5B and C, dark gray bars). In contrast, FCHSD2^S681A^-Myc failed to restore the EGFR trafficking or ERK1/2 sensitivity (Figure 5B and C, black bars); whereas reconstitution with FCHSD2^S681E^-Myc increased rates of EGFR endocytosis and reduced the level of surface EGFR relative to siFCHSD2-treated cells, both of which were resistant to ERK1/2 inhibition (Figure 5B and C, twill bars).

**Figure 5.**
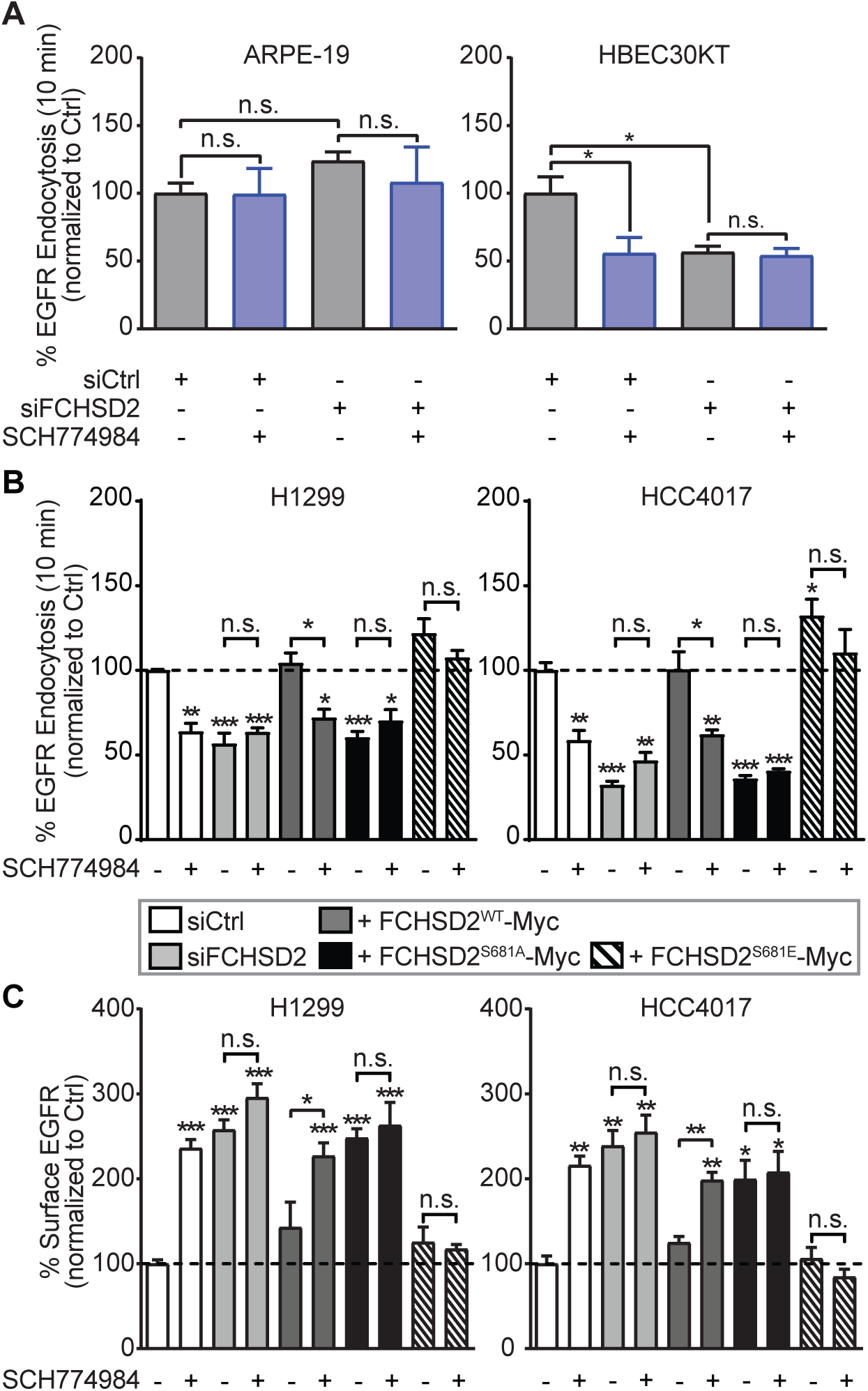
FCHSD2 mediates EGFR trafficking in cancer cells. (*A*) Endocytosis of EGFR was measured in control and FCHSD2 siRNA-treated ARPE-19 and HBEC30KT cells without or with the ERK1/2 inhibitor (SCH772984, 10 µM) treatment. (*B*) Endocytosis of EGFR was measured in the control, FCHSD2 knockdown (KD) cells, or FCHSD2 KD cells reconstituted with FCHSD2^WT^-Myc, FCHSD2^S681A^-Myc, or FCHSD2^S681E^-Myc, in the absence or presence of the ERK1/2 inhibitor (SCH772984, 10 µM). (*C*) Surface levels of EGFR in the cells as described in (*B*). All data were normalized to control and represent mean ± SEM (*n* = 3). Two-tailed Student’s *t* tests were used to assess statistical significance for comparison with Ctrl and for the indicated dataset. n. s., not significant, \**P* < 0. 05, \*\**P* < 0.005, \*\*\**P* < 0.0005.

### FCHSD2 Negatively Regulates EGFR Signaling, Proliferation and Migration

We next measured the downstream consequences of these changes in EGFR endocytosis and surface expression on EGFR signaling and cell behavior. Depletion of FCHSD2 resulted in dramatic increases in several EGF-dependent downstream signaling events, including tyrosine phosphorylation of the EGFR, and activation of Akt and ERK1/2 in both H1299 and HCC4017 cells (Figure 6A and B). Reintroducing FCHSD2^WT^-Myc, but not FCHSD2^S681A^-Myc, again suppressed these signaling activities (Figure 6A and B). Expression of FCHSD2^S681E^-Myc also reduced signaling activities back to control levels (Figure 6A and B). These findings suggest that the crosstalk between ERK1/2 kinases and FCHSD2 contributes to the endocytic trafficking of EGFRs and reciprocally regulates its downstream signaling activities.

**Figure 6.**
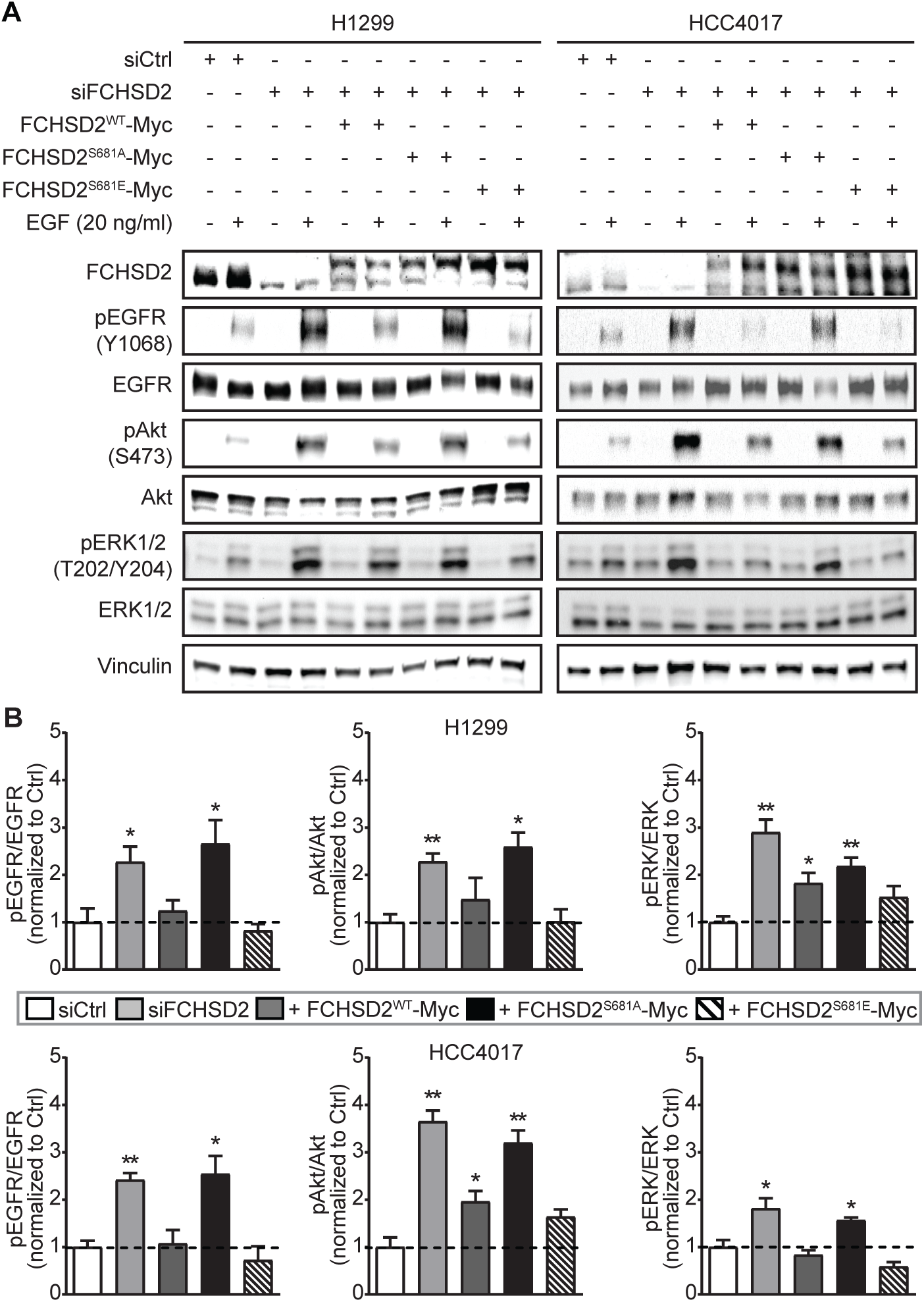
FCHSD2 negatively regulates EGFR signaling in cancer cells. (*A*) Representative western blots of EGFR signaling activities upon EGF stimulation (20 ng/ml) for 10 min after serum starvation for 16 h, in the control, FCHSD2 knockdown (KD) cells, or FCHSD2 KD cells reconstituted with FCHSD2^WT^-Myc, FCHSD2^S681A^-Myc, or FCHSD2^S681E^-Myc. (B) Quantification of pEGFR/EGFR, pAkt/Akt and pERK/ERK intensity ratios in the cells as described in (*A*). All data were normalized to control and represent mean ± SEM (*n* = 3). Two-tailed Student’s *t* tests were used to assess statistical significance for comparison with Ctrl. \**P* < 0.05, \*\**P* < 0.005.

EGFR signaling affects both cancer cell proliferation and migration (44). Therefore, we examined the effects of FCHSD2-dependent alterations in the endocytic trafficking of EGFR and its signaling activities on these cancer cell behaviors. Loss of FCHSD2 increased the rates of proliferation (Figure 7A) in the NSCLC cells. These effects were reversed upon reconstitution with FCHSD2^WT^-Myc or FCHSD2^S681E^-Myc, but less so by the FCHSD2^S681A^-Myc (Figure 7A).

**Figure 7.**
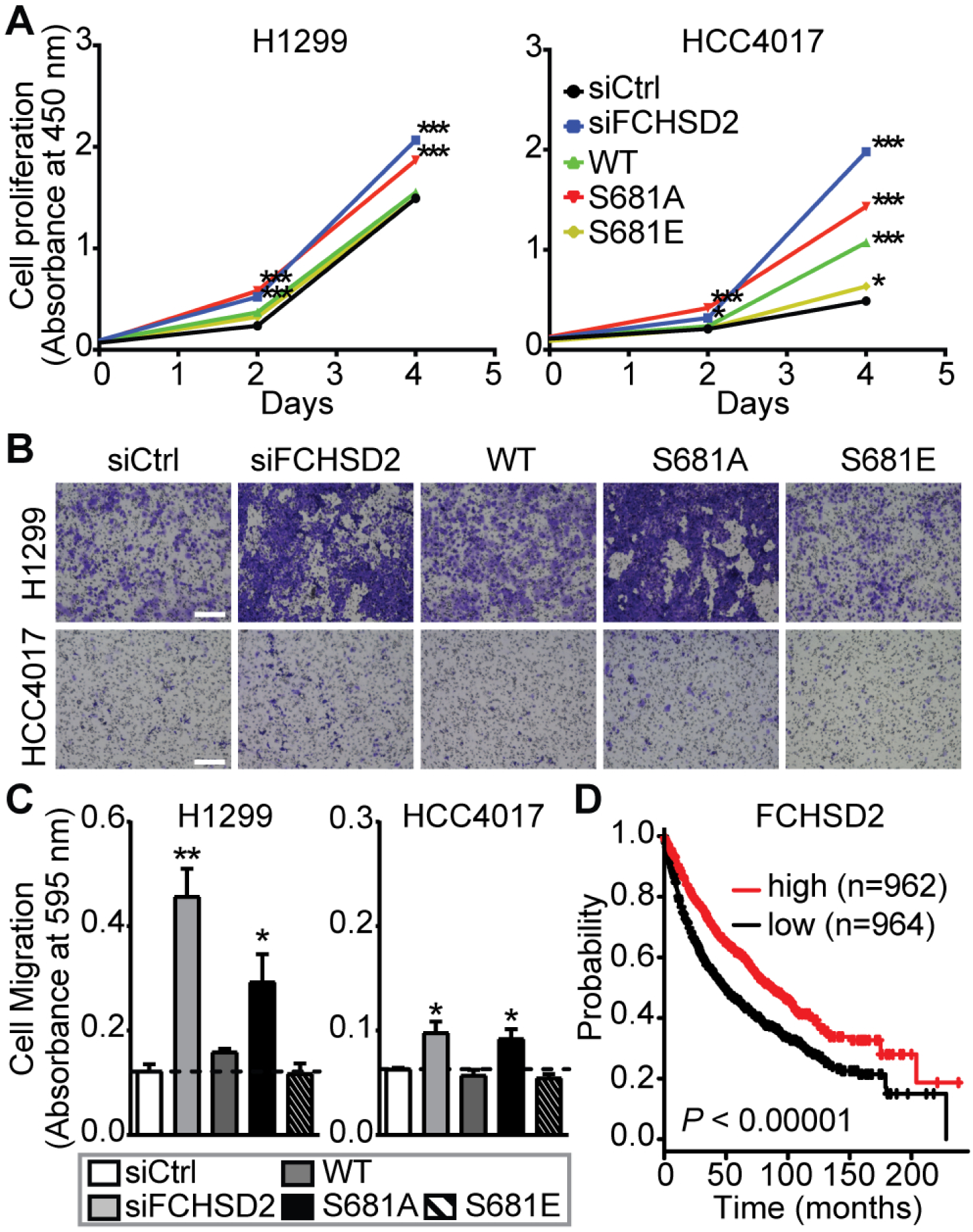
FCHSD2 functions as a negative regulator for cancer progression. (*A*) Cell proliferation abilities were measured in the control, FCHSD2 knockdown (KD) cells, or FCHSD2 KD cells reconstituted with FCHSD2^WT^-Myc, FCHSD2^S681A^-Myc, or FCHSD2^S681E^-Myc. (*B*) Representative images of cell migration in the cells as described in (*A*). Scale bars: 250 µm. (*C*) Quantification of migrating cells in (*B*). Data in (*A*) and (*C*) represent mean ± SEM (*n* = 3). Two-tailed Student’s *t* tests were used to assess statistical significance for comparison with Ctrl. \**P* < 0.05, \*\**P* < 0.005, \*\*\**P* < 0.0005. (D) Kaplan-Meier survival analysis of NSCLC patients was performed in FCHSD2 high- and low-expression cohorts.

Similarly, EGF-dependent migration through transwell filters was enhanced by siRNA knockdown of FCHSD2 (Figure. 7B, C). Reconstitution with either FCHSD2^WT^-Myc or FCHSD2^S681E^-Myc returned migration rates to control levels, whereas reconstitution with FCHSD2^S681A^-Myc less potently suppressed the migration abilities (Figure 7A-C), especially in the more rapidly migrating H1299 cells. and migration (Figure 7B and C)

Finally, we examined the relationship between FCHSD2 expression and survival by mining clinical data and found that patients with relatively high FCHSD2 expression had significantly better survival rates than those in the low-expression group (Figure 7D). Together, these data suggest that FCHSD2 functions as a negative regulator for cancer aggressiveness.

## Discussion

The reciprocal crosstalk between signaling and CME in cancer cells contributes to abnormal trafficking of surface receptors and altered downstream signaling (3, 7, 18). These findings have led to the hypothesis that CME can be ‘adapted’ by cancer cells to support enhanced tumor cell proliferation and metastasis (8). Here we explored the scope of this crosstalk and found that CME can be differentially regulated in cancer cells by several oncogenic signaling kinases. These studies reveal a more extensive crosstalk between signaling and CME than previously appreciated. Further studies are needed to dissect the direct or indirect roles and relevant substrates for the many signaling pathways that selectively impinge on CME in cancer cells.

We focused on the effects of ERK given its importance as a major driver and therapeutic target for multiple cancers (27-30). ERK1/2 inhibition decreased the rates of CME of TfnR and EGFR in cancer cells by specifically inhibiting CCP initiation, the critical first step in CME. ERK1/2 accelerates CME and CCP initiation rates through phosphorylation and activation of FCHSD2. When activated, FCHSD2 negatively regulates EGFR signaling, cell proliferation and migration in NSCLC cells. This study, together with our previous findings (15, 18), strongly supports the concept of adaptive CME and reveals a new mechanism for the reciprocal regulation of signaling and CME in cancer cells.

Although previously not studied in this context, the domain structure of FCHSD2 as well as its known protein interactions (Supplemental Figure 4) are suggestive of a role in CME (47, 48). FCHSD2 is a member of the mild curvature-generating F-BAR family of proteins (49), whose prototypical members, FCHo1/2, have been shown to function in the early stages of CCP initiation and stabilization (41). Here we show that when activated downstream of ERK1/2, FCHSD2 is selectively required for CCP initiation in NSCLC cells. Further studies are needed to probe the underlying mechanisms.

Many F-BAR proteins contain SH3 domains (41, 50) and FCHSD2 encodes two that interact with other components of the endocytic machinery, including dynamin, Wiskott–Aldrich Syndrome protein (WASp) and intersectin1/2 (ITSN1/2) (39, 51). Which of these partners, if any, are required for FCHSD2 function in CCP initiation remains to be determined. In this regard, a recent study, which appeared while preparing this manuscript, reported a role for FCHSD2 in CME using HeLa cells as model (52). These authors showed that FCHSD2 is recruited to CCPs through interactions with intersectin and speculated that it functioned to regulate actin assembly during intermediate stages of CME. While there are some discrepancies between the two studies, with regard to the stage at which FCHSD2 functions, in general the results are complementary and confirm a role for FCHSD2 in CME.

Much more is known about Nervous Wreck (Nwk), the *Drosophila* orthologue of FCHSD2, although it has been studied exclusively in the context of endocytic trafficking and bone morphogenic protein (BMP) signaling at neuromuscular junctions (39, 40, 51). Nwk physically and functionally interacts with components of the endocytic machinery, including WASp and dynamin through its SH3a domain, and Dap160, the *Drosophila* homolog of ITSN1, through its SH3b domain (51). SH3b also binds, albeit in an unconventional manner (40), to the endosomal sorting factor, SNX16. Functional studies link Nwk to sorting of BMP within the endosomal pathway, particularly recycling endosomes (40, 51). Whether FCHSD2 also functions at the recycling endosome in mammalian cells remains to be determined. Interestingly, just as FCHSD2 appears to negatively regulate EGFR signaling and cell proliferation, Nwk is known to negatively regulate BMP signaling, as well as synaptic growth (39, 51).

Nwk displays intramolecular, autoinhibitory interactions between its F-BAR and C-terminal SH3b domain that inhibit both membrane binding of the F-BAR domain and protein interactions with the SH3a domain (53). Release of autoinhibitory SH3b domain interactions is required for activation of Nwk. We found that ERK1/2-mediated phosphorylation of FCHSD2 on S681 is critical for its activity. This phosphorylation site, which resides in the C-terminal region distal to SH3b, is not conserved in Nwk (Supplemental Figure 4) and may reflect an additional level of regulation that renders mammalian FCHSD2 activity responsive to extracellular signaling. We speculate that phosphorylation of S681 either directly disrupts the autoinhibitory interactions of the adjacent SH3b domain or indirectly increases the binding affinity of SH3b domain ligands to relieve the autoinhibition. Alternatively, phosphorylation may alter FCHSD2 interactions with other proteins independently of its own activation. Clearly, further investigation is needed to test these hypotheses regarding the regulation of FCHSD2.

Our results extend FCHSD2 function to outside the nervous system and to earlier stages of the endocytic pathway. Strikingly, FCHSD2 becomes the third component of the neuronal endocytic machinery, along with CLCb and Dyn1 (45), that allows for differential regulation of CME in cancer cells. Endocytic trafficking plays a critical role in regulating and maintaining signaling at the synapse. Whether adaptive endocytosis can result from co-opting other components of the synaptic machinery to alter endocytic trafficking and signaling properties in cancer cells remains to be determined.

Hyperactivity of EGFR signaling at the cell surface is a common feature among different types of cancers and widely considered advantageous for tumor progression (46). Why then, would ERK1/2 activation downstream of EGFR activate a mechanism that enhances CCP initiation and EGFR endocytosis and suppresses EGFR proliferative and migratory signaling? We propose several, not mutually exclusive possibilities for the functional significance of this potentially negative feedback loop. One possibility relates to the well-established spatiotemporal regulation of EGFR signaling (54, 55) and differences in downstream signaling triggered by plasma membrane-associated vs endosome-associated EGFR (56-60). Thus, ERK1/2 and FCSDH2- dependent increases in EGFR uptake, and its redistribution to endosomal compartments could function to alter signaling pathways and downstream cellular responses in ways beneficial to the metastatic cancer cell. A second possibility relates to the documented saturability of EGFR sorting within endosomal compartments (61). Thus, the increased efficiency of EGFR uptake might be needed to ensure saturation of downstream endosomal sorting machinery leading to increased rates of recycling and sustained EGFR signaling. Alternately, buffering of EGFR signaling may be needed within the tumor environment where local concentrations of EGF can be high compared to that seen by metastatic cells (62). Finally, recent evidence has suggested that a subset of EGFR signaling might occur specifically within CCPs (63). Thus, ERK and FCHSD2-dependent increases in CCP initiation might function to increase the numbers of these signaling platforms that trigger a subset of downstream signaling responses. Finally, it’s important to note that both FCHSD2 and its *Ds* homologue, Nwk, function as negative regulators of signaling (this study and (39, 40, 51)). Thus, FCHSD2 activity may play a role in buffering EGFR signaling in normal cells, as we observed in EGF-dependent HBEC30KT cells. In this scenario, its specific upregulation in cancer cells could reflect a compensatory mechanism to counteract the detrimental effects of hyperactive EGFR signaling. Indeed, high levels of FCHSD2 expression correlate with better survival rates in human lung cancer patients.

Overall, our study reveals extensive crosstalk between signaling and clathrin-mediated endocytosis, often specific to cancer cells. Based on our findings reported here and elsewhere (64, 65), we suggest that cancer cells adapt this crosstalk as a determinant for tumor progression. These cancer cell-specific adaptations of the endocytic pathway may provide opportunities of the development of novel therapeutic strategies against cancer.

## Material and Methods

### Cell Culture and Reagents

ARPE-19 cells (from ATCC) were cultivated in DMEM/F12 (Thermo Fisher Scientific) supplemented with 10% (vol/vol) FCS (HyClone). H1299 and HCC4017 NSCLC cells (from John Minna, UT Southwestern Medical Center, Dallas) were grown in RPMI 1640 (Thermo Fisher Scientific) supplemented with 10% (vol/vol) FCS. HBEC30KT, non-transformed human bronchial epithelial cells (from John Minna, UT Southwestern Medical Center, Dallas) were cultivated in complete keratinocyte-SFM medium (Thermo Fisher Scientific). MCF10A cells (from ATCC) were grown in complete mammary epithelial cell growth medium (Lonza). MDA-MB-231 breast adenocarcinoma cells (from R. Brekken, UT Southwestern Medical Center, Dallas) were grown in DMEM containing high glucose medium (Thermo Fisher Scientific), supplemented with 10% (vol/vol) FCS. The following inhibitors and their final concentrations, shown in parenthesis, were used in this study: Akt inhibitor X (10 µM), the CaMK-II inhibitor KN93 (3.7 µM), the CDC42 inhibitor ML 141 (20 µM), the CDK inhibitor Aminopurvalanol A (200 nM), the EPAC inhibitor ESI-09 (32 µM), the ERK inhibitor FR180204 (10 µM), the GSK3β inhibitor CHIR-99021 (10 µM) and Src inhibitor I (440 nM) from MilliporeSigma. The B-Raf inhibitor SB590885 (20 nM), the ERK inhibitor SCH772984 (10 µM), the JAK inhibitor CP 690550 citrate (10 nM) and the JNK inhibitor SP600125 (900 nM) from Selleck Chemicals. The CK2 inhibitor TTP 22 (1 µM), the MLCK inhibitor ML-7 HCl (3 µM), the mTOR inhibitor rapamycin (10 nM), the p38 inhibitor SB239063 (450 nM) and the PKA inhibitor KT-5720 (600 nM) were from Santa Cruz. The MEK inhibitor GSK1120212 (10 µM) and the PI3K inhibitor BKM120 (1 µM) were from MedChemExpress. The PDK1 inhibitor GSK2334470 (100 nM), the PTEN inhibitor VO-OHpic (460 nM) and the ROCK inhibitor GSK269962 (40 nM) were from Tocris Bioscience. The PKC inhibitor Gö-6983 (600 nM) was from Cayman Chemical. Protease and phosphatase inhibitor cocktails were from Roche, all other chemicals were reagent grade and purchased from Sigma.

### Endocytosis Assays

TfnR and EGFR endocytosis assays were performed exactly as described (18) using anti-TfnR mAb (HTR-D65) or biotinylated-EGF respectively using cells grown overnight in gelatin-coated 96-well plates and plated at a density of 3×10^4^ cells/well. For TfnR and EGFR endocytosis assays using inhibitors, cells were preincubated in the absence (i.e., control) or presence of the indicated inhibitors and concentrations described above for 30 min at 37°C before internalization assays were performed in the continued absence or in the presence of the respective inhibitors.

### Western Blotting

Cells were washed three times with PBS and harvested/resuspended in 150–200 µl of reducing Laemmli sample buffer. The cell lysate was boiled for 10 min and 50 µg of cell lysate was loaded onto an SDS gel. After transferring to a nitrocellulose membrane. Membranes were blocked with either 5% powdered milk or 5% BSA and probed with antibodies against the following proteins: pERK1/2 T202/Y204 (#4370S, Cell Signaling), ERK1/2 (#4695S, Cell Signaling), pEGFR Y1068 (#3777S, Cell Signaling), EGFR (#4267S, Cell Signaling), pAkt S473 (#4060L, Cell Signaling), Akt (#9272S, Cell Signaling) and Vinculin (#V9131, MilliporeSigma), which was used as internal controls, according to the manufacturers’ instructions. For FCHSD2 we used #PA5-58432 (Thermo Fisher Scientific), which detected an ∼85 kD band that was specifically knocked down by siFCHSD2 treatment at 1:500 dilution. Note that the manufacturer’s data shows a 100 kD band, which we also detected, but this band did not respond to siRNA knockdown. The calculated MW for the 740aa variant of FCHSD2 (Accession number O94868) is ∼84 kD. Horseradish peroxidase (HRP)-conjugated secondary antibodies (#G21234 and # G21040, Thermo Fisher Scientific) were used according to the manufacturers’ instructions.

Quantitative analysis was performed by using ImageJ software (NIH). For EGF-dependent signaling, the H1299 and HCC4017 cells (5×10^5^ cells) were seeded in 6-well plates containing RPMI 1640 with 10% FCS. Eight hours after seeding, cells were washed three times with PBS and starved in RPMI 1640 without FCS for 16 h. The cells then were untreated or treated with 20 ng/ml of EGF for 10 min. After the EGF treatment, cells were washed three times with PBS and harvested/resuspended in 150–200 µl of reducing Laemmli sample buffer, and the cell lysates were subjected to Western blotting as described above.

### TIR-FM and Image Data Analysis

Total internal reflection fluorescence microscopy (TIR-FM) was performed as previously described (13). Briefly, ARPE-19 and H1299 stably expressing eGFP-CLCa and HCC4017 expressing SNAP-CLCa were imaged using a 100x 1.49 NA Apo TIRF objective (Nikon) mounted on a Ti-Eclipse inverted microscope with Perfect Focus System (Nikon). TIR-FM illumination was achieved using a Diskovery Platform (Andor Technology). During imaging, cells were maintained at 37°C in complete culture medium. Time-lapse image sequences were acquired at a penetration depth of 80 nm and a frame rate of 1 Hz using a sCMOS camera with 6.5 µm pixel size (pco.edge). For TIR-FM using inhibitors, cells were preincubated in the absence (i.e., control) or presence of the indicated inhibitors described above for 30 min at 37°C, followed by imaging in the absence or in the presence of the respective inhibitors. CCP lifetime distribution and initiation density analyses were carried out in Matlab (Math-Works, Natick, MA, USA), using cmeAnalysisPackage (66).

### Immunoprecipitation, Mass Spectrometry and Proteomic Analysis

HCC4017 stably expressing FCHSD2-Myc were washed three times with PBS and starved in RPMI 1640 without FCS for 16 h. The cells then were untreated or treated with 100 ng/ml of EGF for 10 min. After the EGF treatment, cells were washed three times with PBS and lysed for 30 min at 4°C in lysis buffer (20 mM HEPES (pH 7.4), 150 mM KCl, 2 mM MgCl_2_, 1 mM Na_3_VO_4_, 1X Protease Inhibitor Cocktail, 1X Phosphatase Inhibitor Cocktail) containing 0.2% Triton X-100. Lysates were clarified at 12,000g for 15 min at 4°C. After quantification using BCA method (Thermo Fisher Scientific), about 3 mg of total protein lysate in 500 µl lysis buffer was used for each immunoprecipitation. Anti-Myc-DDK antibody (#TA50011-100, OriGene) was used to immunoprecipitate FCHSD2-Myc by incubation with the lysate for 1 h at 4°C. Addition of Protein G beads (approximately 40 µl) (MilliporeSigma) in the lysate followed by gentle rotation for 1 h at 4°C allowed binding of target proteins. Beads were washed three times in lysis buffer and then denatured using reducing Laemmli sample buffer, boiled and run on SDS-PAGE gel.

Protein gel pieces was reduced and alkylated with DTT (20 mM) and iodoacetamide (27.5 mM). A 0.01 µg/µl solution of trypsin in 50 mM triethylammonium bicarbonate (TEAB) was added to completely cover the gel, allowed to sit on ice, and then 50 µl of 50 mM TEAB was added and the gel pieces were digested overnight (Pierce). Following solid-phase extraction cleanup with an Oasis HLB µelution plate (Waters), the resulting peptides were reconstituted in 10 ul of 2% (v/v) acetonitrile (ACN) and 0.1% trifluoroacetic acid in water. Five ul of this were injected onto an Orbitrap Fusion Lumos mass spectrometer (Thermo Electron) coupled to an Ultimate 3000 RSLC-Nano liquid chromatography systems (Dionex). Samples were injected onto a 75 µm i.d., 50-cm long EasySpray column (Thermo), and eluted with a gradient from 1-28% buffer B over 60 min. Buffer A contained 2% (v/v) ACN and 0.1% formic acid in water, and buffer B contained 80% (v/v) ACN, 10% (v/v) trifluoroethanol, and 0.1% formic acid in water. The mass spectrometer operated in positive ion mode with a source voltage of 2.4 kV and an ion transfer tube temperature of 275 °C. MS scans were acquired at 120,000 resolution in the Orbitrap and up to 10 MS/MS spectra were obtained in the Orbitrap for each full spectrum acquired using higher-energy collisional dissociation (HCD) for ions with charges 2-7. Dynamic exclusion was set for 25 s after an ion was selected for fragmentation.

Raw MS data files were analyzed using Proteome Discoverer v2.2 (Thermo), with peptide identification performed using Sequest HT. Fragment and precursor tolerances of 10 ppm and 0.6 Da were specified, and three missed cleavages were allowed. Carbamidomethylation of Cys was set as a fixed modification. Oxidation of Met and phosphorylation of Ser, Thr, and Tyr were also set as variable modifications and were localized using ptmRS.

### Mutagenesis of FCHSD2 and generation of phosphomutants

The FCHSD2 (NM_014824) Human Tagged ORF clone lentiviral construct was purchased from Origene (#RC221241L3). The FCHSD2-Myc tagged phosphomutants (S681A, S681E) were generated by standard site directed mutagenesis (67). The primer sequences for the mutagenesis of S681A and S681E are 5’- CTGTACTTTCCCCGG<u>G</u>CTCCTTCAGCAAACG-3’ and 5’- GCTCCCTGTACTTTCCCCGG<u>GAG</u>CCTTCAGCAAACGAAAAAAG-3’, respectively. All primers were purchased from Integrated DNA Technologies.

### siRNA Transfection and Reconstitution with FCHSD2-Myc Constructs

Cells were treated with the siRNA pool targeting FCHSD2 (#E-021240-00-0010, Dharmacon) using RNAiMAX (Thermo Fisher Scientific) to silence the endogenous protein. Briefly, 50 nM of the indicated siRNA pool and 6.5 µl of RNAiMAX reagent were added in 1 ml of OptiMEM (Thermo Fisher Scientific) in each well of a 6-well plate and incubated for 20 min at room temperature. Cells were resuspended in 1 ml of culture medium, seeded in each well of a 6- well plate containing the mixed siRNA-lipid complex and incubated for 48 h, followed by experiments. The AllStars Negative siRNA non-targeting sequence was purchased from Qiagen (#SI03650318).

Reconstitution with FCHSD2^WT^-Myc, FCHSD2^S681A^-Myc, or FCHSD2^S681E^-Myc fusion proteins in H1299 and HCC4017 FCHSD2 siRNA-mediated knockdown cells was performed using lentiviral vectors. The lentiviral vectors and lentiviral packing plasmids (pSPAX2 (Gag/Pol) and pMD2G) were co-transfected into HEK293T cells using Lipofectamine 2000 (Thermo Fisher Scientific) according to manufacturer’s instructions for virus production. The virus conditioned media was used to infect target cells in the presence of 10 µg/ml polybrene (MilliporeSigma). Cells were collected 48 h post-infection for experiments.

### Cell Proliferation

Cells were seeded on 96-well plates at a density of 3×10^3^ cells/well and incubated for the indicated time points. The cell proliferation ability was measured by using the CCK-8 Counting Kit (Dojindo Molecular Technologies) according to the manufacturer’s instructions. Briefly, cells were incubated with culture medium containing the CCK-8 solution for 1 h at 37°C, and then the absorbance was read at 450 nm.

### Cell Migration

Tumor cell migration ability was evaluated in 6.5 mm Transwell with 8.0 µm pore polyester membrane insert (#3464, Corning) in 24-well plates. 1×10^5^ cells were resuspended in 300 µl of RPMI 1640 without FCS and seeded on each transwell insert, and each well of the 24-well plate was filled with 700 µL of complete culture medium. After incubation for 16 h at 37 °C, the cells remaining on the upper surfaces of the membrane were removed by wet cotton swabs. Cells that had migrated to the lower surfaces of the membrane were fixed with ice-cold methanol (Pharmco) for 10 min and stained with 0.5% crystal violet (MilliporeSigma) for 15 minutes. After imaging, cells were washed three times with ddH_2_O and incubated in 1% SDS solution (MilliporeSigma) for 5 min at room temperature and the number of cells quantified by the absorbance at 595 nm.

### Analysis of Kaplan-Meier Survival Data

NSCLC patient survival data was downloaded from the Kaplan Meier plotter database (68). Analysis of NSCLC patients was performed in FCHSD2 high and low expression cohort. *P* value was calculated by logrank test (68).

## Acknowledgments

We are grateful to Richard Carr III, for initiating this direction of research while he was a postdoctoral fellow in the Schmid lab. We thank members of the Schmid lab for critically reading the manuscript and Heather Grossman, Kim Reed and Marcel Mettlen for technical assistance in plasmid preparation and microscopy, respectively. We thank Andrew Lemoff and the UTSW Proteomics Core Facility for help with mass spectrometry and proteomic analysis. The work was supported by NIH grants GM73165 to SLS and Gaudenz Danuser, and MH61345 to SLS.

## Author contributions

G-Y.X and S.L.S designed the research; G-Y.X. performed the research, analyzed data and prepared the figures; A.M. prepared FCHSD2 phosphomutant constructs and lentiviral vectors. G-Y.X. and S.L.S. wrote the paper. All authors read and commented on the MS.

---

**Figure S1.**
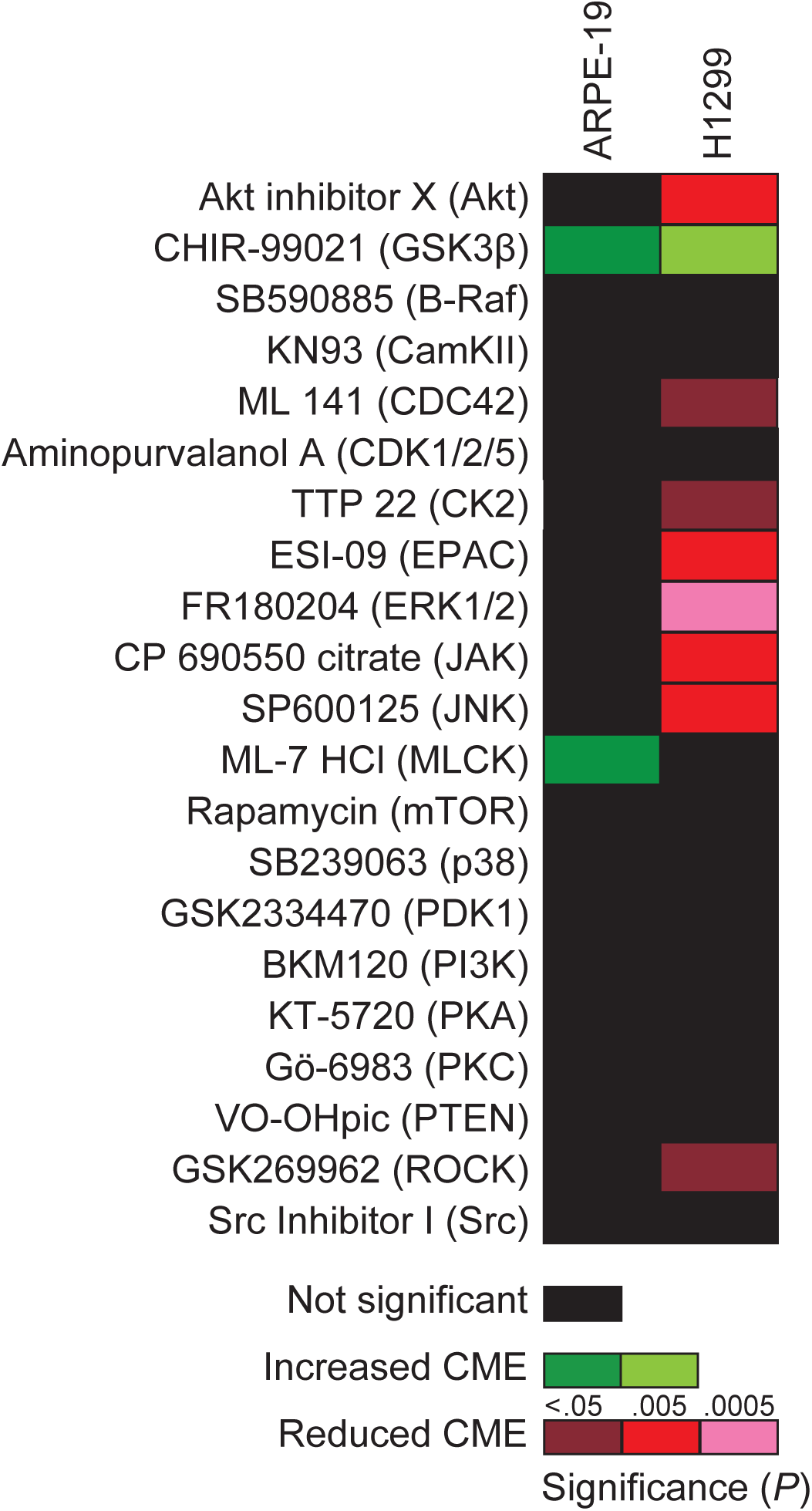
Analysis of the effects of kinase inhibitors on the endocytosis of TfnR in human non-cancerous ARPE-19 and H1299 NSCLC cells. Heatmap illustrating the effects of the indicated inhibitors on TfnR endocytosis in non-cancerous ARPE-19 cells and H1299 NSCLC cells.

**Figure S2.**
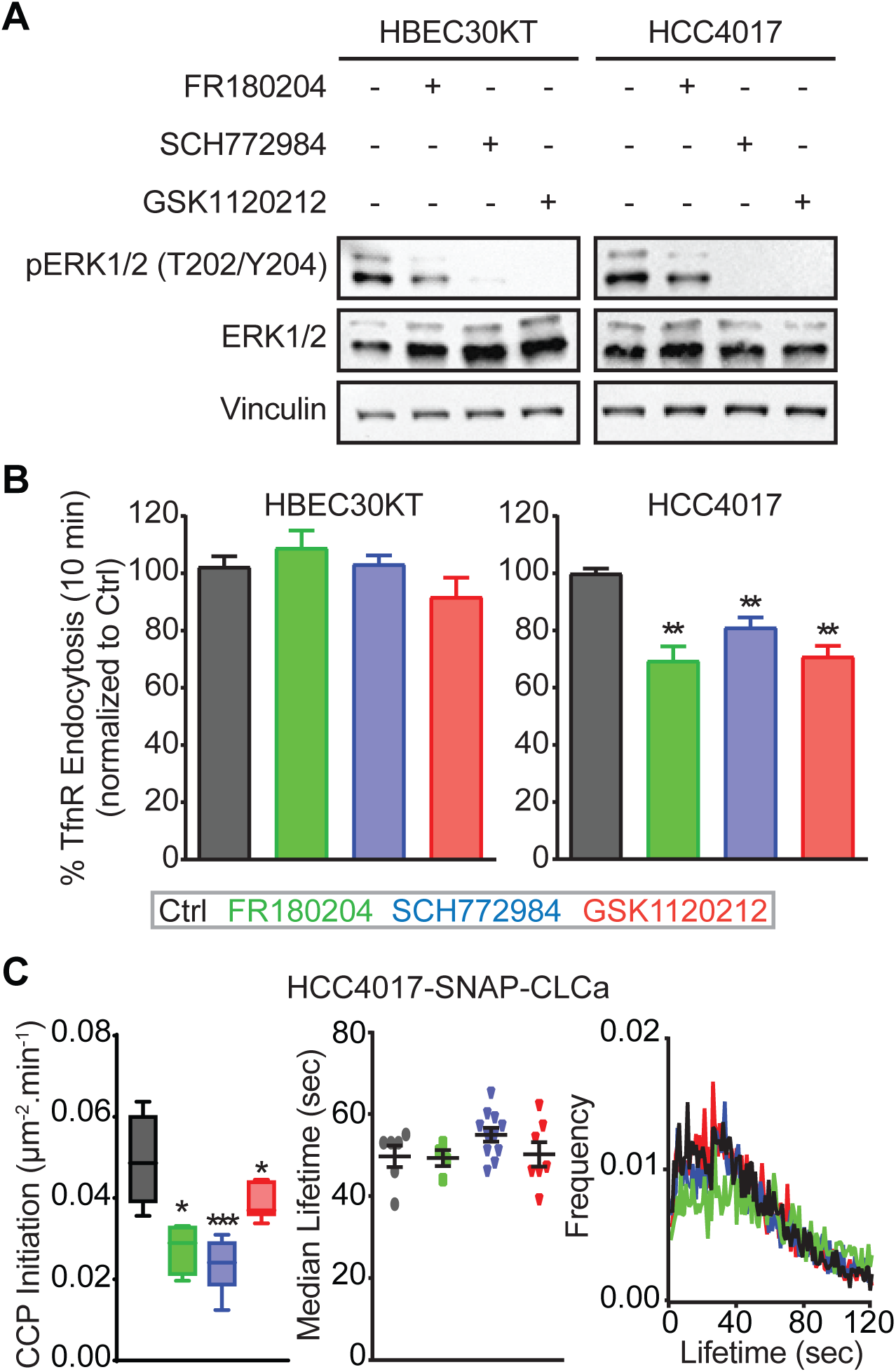
ERK1/2 specifically affects CME activities in cancer cells. (*A*) Representative western blots to measure the efficiencies of kinase inhibitors in reducing ERK1/2 phosphorylation in control, ERK1/2 inhibitor (FR180204 and SCH772984, 10 µM) and MEK1/2 inhibitor (GSK1120212, 10 µM) treated HBEC30KT and HCC4017 cells. (*B*) Endocytosis of TfnR was measured in control and the inhibitor, as described in (*A*), -treated HBEC30KT and HCC4017 cells. All data were normalized to control and represent mean ± SEM (*n* = 3). (*C*) Average lifetime distributions (*left*), and rates of initiation (# CCP/µm^2^/min, *right*) of *bona fide* CCPs in control and the inhibitor, as described in (*A*), -treated HCC4017 cells. Data were obtained from at least 15 cells/condition, the box plots represent median, 25th and 75th percentiles, and outermost data points. Two-tailed Student’s *t* tests were used to assess statistical significance for comparison with Ctrl. \**P* < 0. 05, \*\**P* < 0.005, \*\*\**P* < 0. 0005, n. s., not significant.

**Figure S3.**
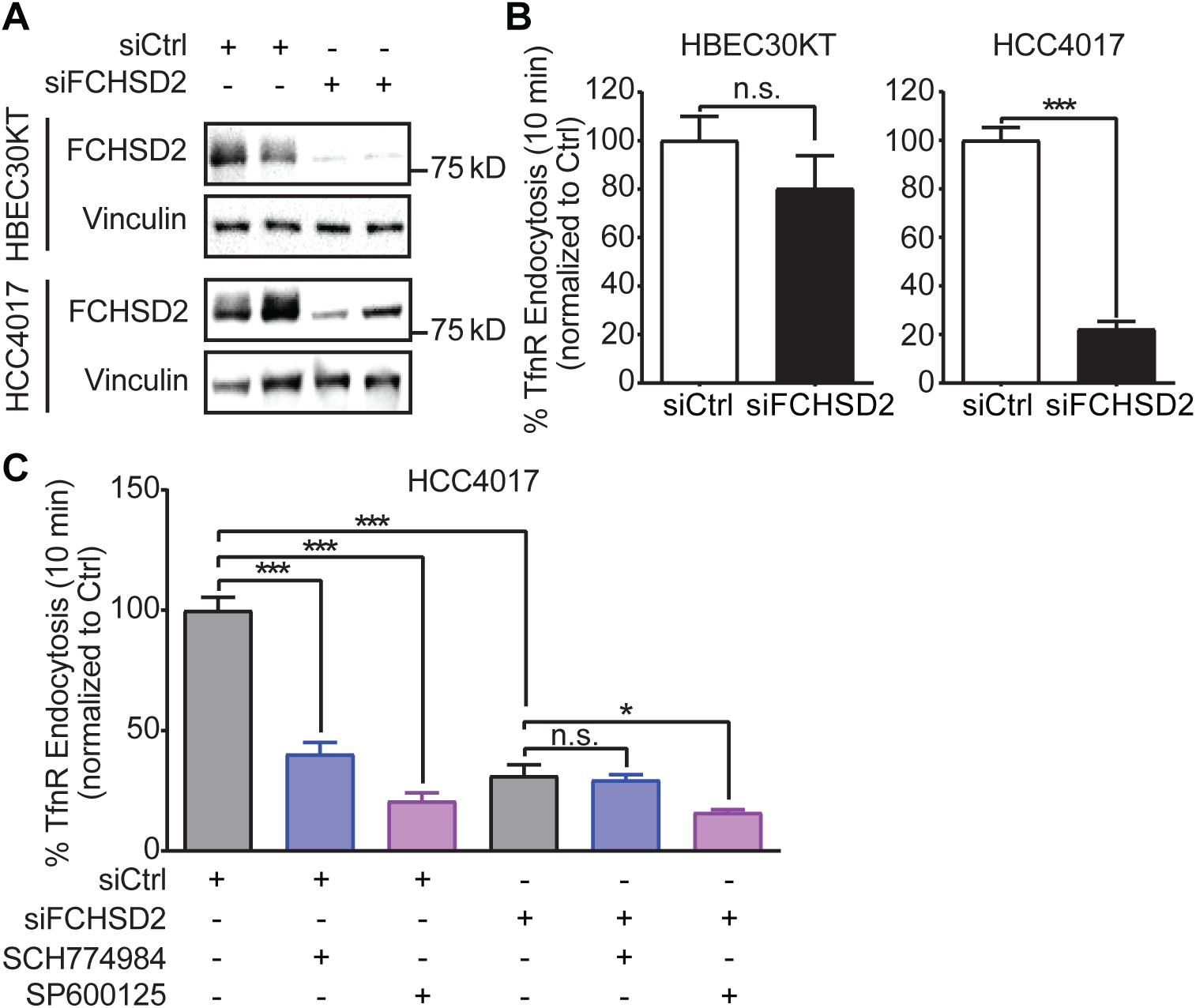
FCHSD2 is specifically required for CME in cancer cells. (*A*) Representative western blots and the quantification of FCHSD2 knockdown efficiency in control and FCHSD2 siRNA-treated HBEC30KT and HCC4017 cells. (*B*) Endocytosis of TfnR was measured in control and FCHSD2 siRNA-treated HBEC30KT and HCC4017 cells. All data were normalized to control. (*C*) Endocytosis of TfnR was measured, as described in (*B*), in control and FCHSD2 siRNA-treated HCC4017 cells without or with the ERK1/2 inhibitor (SCH772984, 10 µM) or the JNK inhibitor (SP600125, 10 µM) treatment. All data represent mean ± SEM (*n* = 3). Two-tailed Student’s *t* tests were used to assess statistical significance. n.s., not significant, \**P* < 0.05, \*\*\**P* < 0.0005.

**Figure S4.**
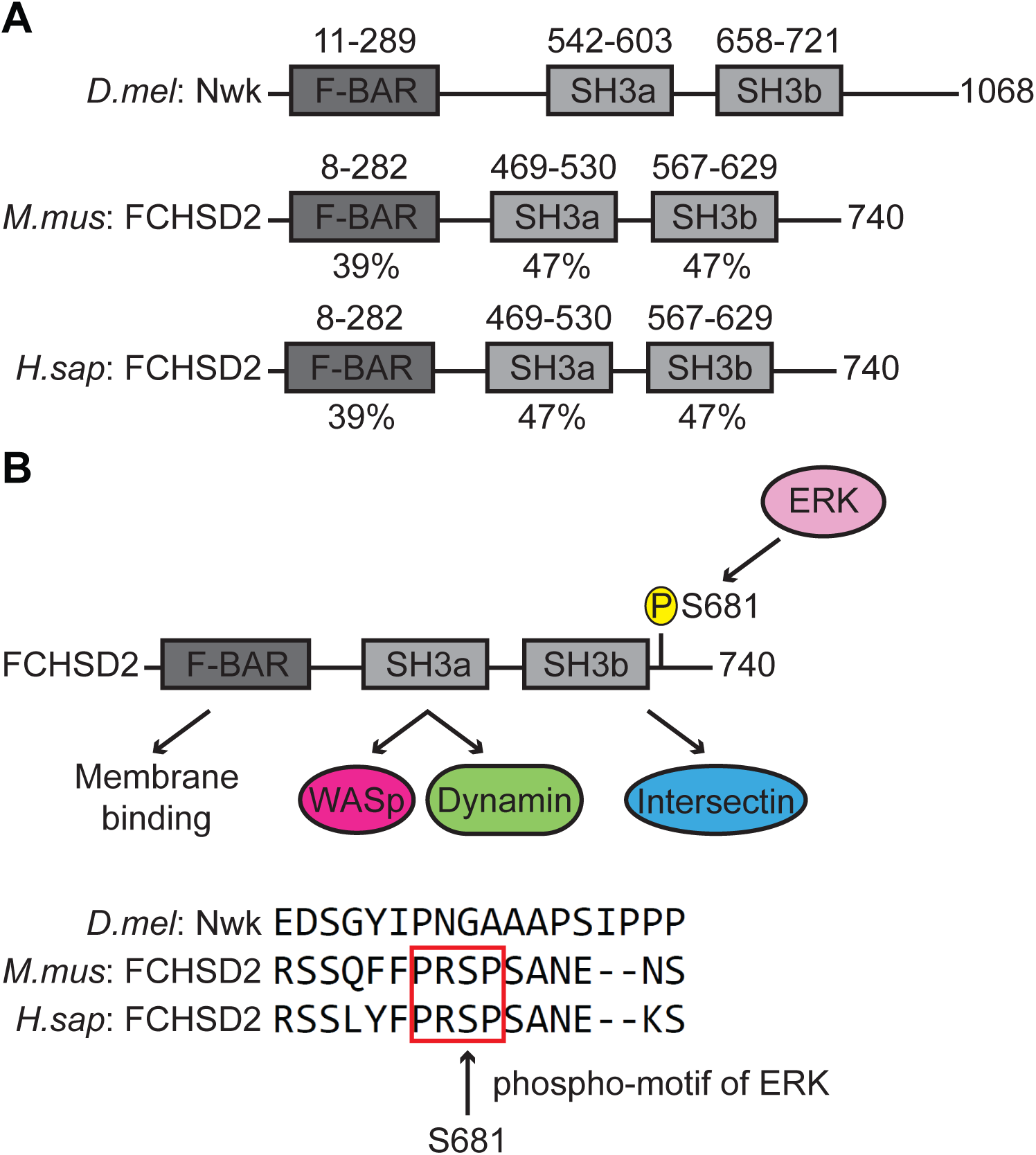
Mammalian FCHSD2 is homologous to fly Nwk and phosphorylated by ERK1/2 kinases. (*A*) Schematic drawing of the domain structures of *Drosophila melanogaster* Nwk, *Mus musculus* and *Homo sapiens* FCHSD2. (*B*) Model of the ERK-mediated phosphorylation site in FCHSD2 and the interactions between FCHSD2, WASp, dynamin, and intersectin. The phosphorylation site in mammalian FCHSD2 is not conserved in Nwk.

